# Ongoing neural oscillations predict the post-stimulus outcome of closed loop auditory stimulation during slow-wave sleep

**DOI:** 10.1101/2021.05.06.443016

**Authors:** Miguel Navarrete, Steven Arthur, Matthias S. Treder, Penelope A. Lewis

## Abstract

The large slow oscillation (SO, 0.5-2Hz) that characterises slow-wave sleep is crucial to memory consolidation and other physiological functions. Manipulating slow oscillations can enhance sleep and memory, as well as benefitting the immune system. Closed-loop auditory stimulation (CLAS) has been demonstrated to increase the SO amplitude and to boost fast sleep spindle activity (11-16Hz). Nevertheless, not all such stimuli are effective in evoking SOs, even if they are precisely phase-locked. Here, we studied whether it is possible to use ongoing activity patterns to determine which oscillations to stimulate in order to effectively enhance SOs or SO-locked spindle activity. To this end, we trained classifiers using the morphological characteristics of the ongoing SO, as measured by electroencephalography (EEG), to predict whether stimulation would lead to a benefit in terms of the resulting SO and spindle amplitude. Separate classifiers were trained using trials from spontaneous control and stimulated datasets, and we evaluated their performance by applying them to held-out data both within and across conditions. We were able to predict both when large SOs will occur spontaneously, and whether a phase-locked auditory click will effectively enlarge them with an accuracy of ~70%. We were also able to predict when stimulation would elicit spindle activity with an accuracy of ~60%. Finally, we evaluate the importance of the various SO features used to make these predictions. Our results offer new insight into SO and spindle dynamics and provide a new method for online optimisation of stimulation.

**HIGHLIGHTS:**

- Random forest classifiers can predict spontaneous and stimulated SOs and spindle amplitudes.
- Morphological wave features predicted the response of SOs and spindles to CLAS.
- SO amplitude during the click is the main predictor for post-stimulus SO amplitude.
- Prediction of spindle activity did not differ in accuracy for stimulated vs spontaneous data.

## 1. INTRODUCTION

Slow wave sleep (SWS) is important for memory consolidation and crucial for metabolic regulation and neural recovery (Klinzing et al., 2019; Nedergaard et al., 2013). This sleep state is characterized by slow wave activity (SWA), mainly associated with continuous epochs of high amplitude slow oscillations (SO, 0.5-2 Hz) (Iber et al., 2007). SOs are thought to be critical for memory consolidation because they provide the necessary neurophysiological conditions for hippocampal-cortical binding (Jiang et al., 2019; Klinzing et al., 2019). Evidence suggest that the coupling of SOs with sleep fast spindles (12-16Hz) provides the ideal timing for transfer of information from the hippocampus to the cortex (Helfrich et al., 2019; Maingret et al., 2016; Peyrache et al., 2009). Therefore, several non-invasive methods have been proposed for boosting SO amplitudes. These have been demonstrated to improve cognitive and physiological characteristics related to this slow wave activity (Marshall et al., 2006; Massimini et al., 2007; Ngo et al., 2013).

Among techniques proposed for increasing the effect of SOs, closed loop acoustic stimulation (CLAS) has proved to be promising in both young (Ngo et al., 2013; Ong et al., 2018) and older participants (Papalambros et al., 2017; Schneider et al., 2020). In CLAS, auditory clicks are applied on the peaks of SOs, increasing SO amplitude and associated phase-locked spindles (Ngo et al., 2013). CLAS has been shown to improve post-sleep memory retention (Ngo et al., 2013; Papalambros et al., 2017) and to positively impact the immune and the autonomic function of sleep (Besedovsky et al., 2017; Grimaldi et al., 2019). Nevertheless, this technique is not without limitations. Previous studies showed that CLAS is a self-limited process, which increases neither SWA across the night, nor the density of slow waves during SWS (Ngo et al., 2015). Likewise, CLAS has limited benefits in terms of improving the consolidation of visual and procedural memories previously associated with SOs (Leminen et al., 2017). Furthermore, increases in SO amplitude and power through this technique are apparently not beneficial enough to increase memory consolidation (Henin et al., 2019), and sensitivity to CLAS stimulation reduces with age (Schneider et al., 2020).

All this prior evidence suggests that the impact of CLAS is not exclusively determined by the effects of the sensory input, but is also influenced by ongoing neural processes in the SO (Navarrete et al., 2020a). Such ongoing processes therefore determine whether an auditory stimulus can increase SO amplitude. In theory, two principal neural processes could be determining the outcome of each auditory click on the SO. The first process relates to the type of wave that is stimulated. For instance, a recent study in rodents suggests dissociable functional roles for fast and slow oscillatory elements during SWA (Kim et al., 2019). Likewise, in humans, different memory mechanisms have been proposed for SO elements depending on their origin, extension and spindle locking (Bernardi et al., 2018; Helfrich et al., 2019; Siclari et al., 2014). Nevertheless, the auditory modulation of SOs by auditory clicks has typically been evaluated as if all waves originated from the same mechanisms, and as if every stimulus were equally efficient for increasing SO function. A second process may be established by the ongoing neurophysiological processes co-occurring with the SO when the stimulus is applied. Previous work theorised that the outcome of the stimulus (measured as ‘no response’, ‘increased SO amplitude’ or ‘sleep arousal’) is determined by a sweet spot in stimulation that, if targeted appropriately, allows boosting of the SO without causing arousal (Bellesi et al., 2014). This work suggested that this sweet spot is determined by the SOs themselves, since SOs can supress inputs from the locus coeruleus (LC), thus preventing arousals in response to sound stimuli while increasing the SO amplitude. Unfortunately, the nature of the neural threshold which determines this sweet spot is still unknown, leaving a lot of uncertainty about the outcome of CLAS stimulation, as little is known about the responsiveness of each individual SO. Currently, there are no a-priory rules to determine when a SO will have a large amplitude spontaneously or when the optimal large-wave response will occur in response to the CLAS click.

In this study, we aimed to examine the factors predicting the spontaneous and evoked response to an auditory click on each SO. We hypothesize that if cortical activity modulates the response of each stimulus by differentiated SO processes, then it would be possible to predict subsequent spontaneous and induced cortical dynamics. To test this, we trained machine learning models to predict the outcome of cortical activity in both spontaneous and stimulation conditions. We used a series of features based on morphological characteristics of the ongoing SO and the estimated timing of the click stimulation. Our results show that the trained models can predict whether subsequent SO amplitude will be high or low relative to the average, and whether an increment of spindle activity will occur. We did this in both unstimulated SWS and CLAS conditions. Furthermore, using a feature importance analysis we then evaluated the variables that allowed us to determine the sensitivity of the ongoing wave to the auditory stimulus. We found that ongoing SWS dynamics predict the event amplitude of both spontaneous and stimulated SOs, and that the threshold of maximal responsiveness is determined by the level of cortical activation during the click as measured by the amplitude of large SOs. We argue that the response to stimulation is mediated by the high drive for cortico-thalamic activation and the reduced cortico-coerulear drive of large SO positive cycles. Our findings support the idea that SWS is comprised of several types of SOs. This analysis of the response to CLAS may help to unravel the function of the SO, as well as to pave the way for more targeted stimulation and enhancement of this key sleep stage.

## 2.1 MATERIALS AND METHODS

### 2.1 Datasets and experimental procedures

The data were taken from (Navarrete et al., 2020a). Briefly, polysomnographic data including EEG and hypnogram of 21 individuals (14 females and mean ± SD age = 25.7 ± 4.7 years). The participants spent two experimental nights in the laboratory undergoing one experimental stimulation (STIM) and one no-stimulation condition (SHAM). The order of experimental conditions was balanced across subjects and separated by at least one week. All protocols were approved by the appropriate ethic committees of local institutions (University of Lübeck and University of Los Andes) and written consents were obtained for each participant.

Acoustic stimuli for the STIM condition consisted of stereophonic clicks of pink noise (50ms duration) with rising and falling slopes (5ms duration), and stimulation timestamps were recorded online when clicks were applied in STIM (stim-click) or for when they were predicted in SHAM (sham-click). For the SHAM condition, the detection protocol was identical to STIM, but the sound was muted. The streamed signal was filtered in the SO frequency band and negative EEG deflections that surpassed an adaptative threshold were identified as a SO down-state during SWS. After each trial, there was a pause for 2.5s before trough detection was resumed. Stimulation was applied during sustained non-rapid eye movement sleep (NREM), including N2 and N3 stages, and this was manually halted if there was visual evidence of arousals or REM.

### 2.2 Sleep and EEG analysis

For the analysis and pre-processing of sleep EEG, we closely followed the procedures from (Navarrete et al., 2020a). Sleep scoring was performed according to ASSM scoring criteria (Iber et al., 2007) by two trained experimenters blinded to stimulation conditions. All artefacts and arousals were marked in the hypnogram. We focused on events detected in Fz for SOs and sleep spindles activity (SA) because these are the locations where SO are more pronounced (Iber et al., 2007). The raw data were resampled at 200Hz with linear interpolation after applying an antialiasing low pass FIR filter. After processing, only stimulations marked during N3 sleep stage were retained. SWA was obtained from the EEG signals filtered between 0.5 - 2 Hz using a zero-phase windowed equiripple FIR filter (3dB at 0.25 and 3.08 Hz; >37 dB at f < 0.01 Hz and f > 4 Hz). Waves were only considered as SOs when their negative deflection had consecutive zero crossings between 0.25 to 1.0 seconds, regardless of the wave amplitude (Riedner et al., 2007). Likewise, spindle activity was determined by applying a zero-phase bandpass FIR filter between 11 and 16 Hz (3 dB at 10.62 and 17.38 Hz; >40 dB at f < 10.01 Hz and f > 18 Hz). Then, the root mean squared (RMS) was computed by using a time window of 0.2 seconds. Rejection criteria included trials where stimulus was applied less than 2 sec apart from any arousal, artefact, or changes to another sleep stage other than N3. For spindle event detection, the candidate events were first detected as discrete events where the RMS signal surpassed a threshold established as the 86.64 percentile (equivalent to 1.5 SD over the mean for a Gaussian distribution) of the spindle activity during N3. Then, spindles were identified as events with duration between 0.3 and 3 seconds (Warby et al., 2014), with at least five oscillations, a unimodal peak in the spindle frequency band and decreasing power for higher frequencies computed by the Morlet wavelet (Purcell et al., 2017). The frequency band for fast spindles (FS) was defined as 11 – 16 Hz, and in the 9 – 12Hz for slow spindles (SS). Finally, to avoid differences due to phase distribution, all ERP analyses were computed for events where sham and stim click were applied within the phase interval Φ: −π/4 to π/4 (Navarrete et al., 2020a).

### 2.3 Selection of features and labels for classification

For the selection of the model features, we selected a series of morphological wave features taken before and during the stimulation of the ongoing wave as shown in Figure 1. These predictor features, further defined in Table 1, describe morphological characteristics of the pre-stimulus slow-wave structure (*v.Neg, t.Neg, t.PosNeg, areaPos, waveRatio* and *slopeRatio* in table 1), structural characteristics of SO detection and peak estimation (*cosPhase*, *sinPhase*, *v.Stim* and *t.Stim* in table 1) and long term dynamics of spindle activity (*FSonStim*, *SSonStim*, *FS_Lag* and *SS_Lag* in table 1). Additionally, to control for random effects in the analysis of feature importance, two dummy variables were included. These were the features representing the subject ID and a random generated number (*sbjID* and *random* in table 1).

**FIGURE 1.**
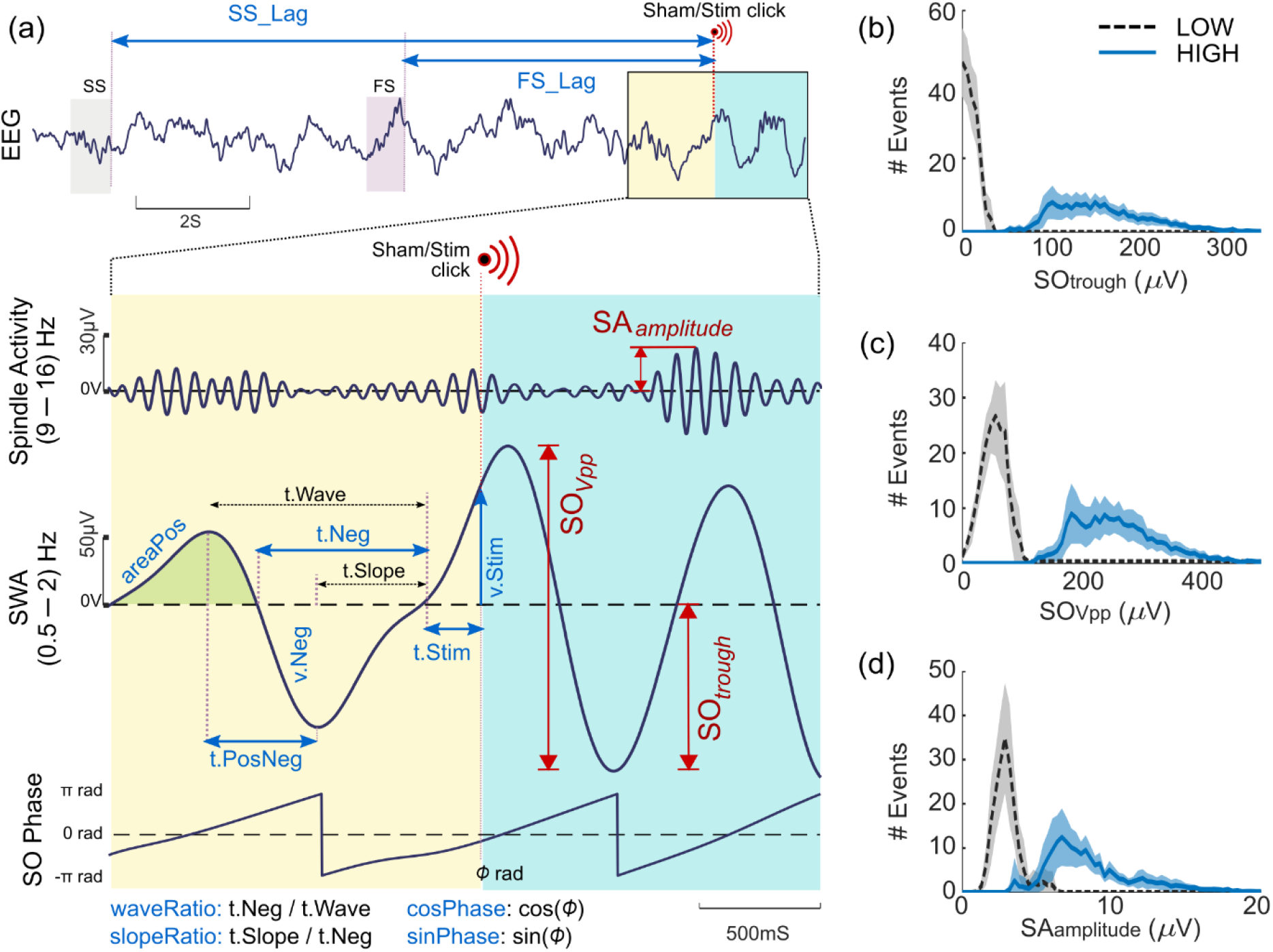
Selection of features for training of classifiers and the post-stimulus measures evaluated as response to click stimulation together with between-subject average distribution of the events labelled LOW and HIGH. (a) Four post stimulus measures were assessed in the study (in red; SO_*Vpp*_: peak-to-peak SO amplitude, SO_*trough*_: SO’s trough amplitude and SA_*amplitude*_: spindle activity amplitude). The features used to train the machine learning model are marked in blue, and they include information from pre-stimulus fast spindles (FS: 12 – 16Hz) or slow spindles (SS: 9 – 12Hz) (e.g. FS_Lag: Time to the previous FS, SS_Lag: Time to the previous SS), as well as information of the pre-stimulus SO wave (e.g. v.Neg: negative voltage, t.Neg: time of negative wave) and information about the timing of stimulus (e.g. v.Stim: SO voltage during stimulation, sinPhase: sinus of phase of stimulation). At the stimulus time (Stim) an auditory click was applied in STIM trials whereas a muted sham-click was marked in SHAM trials. A meticulous description of all evaluated features is presented in Table 1. (b) Between-subject distribution of LOW and HIGH classes for SO trough amplitudes (SO_*trough*_). (c) Between-subject distribution of LOW and HIGH classes for SO peak-to-peak amplitudes (SO_Vpp_). (d) Between-subject distribution for LOW and HIGH classes for spindle activity amplitude (SA_*amplitude*_). Dispersion areas represent mean ± 95% CI of the distribution.

**TABLE 1.** Selection of wave and stimulation characteristics used as predictors for classification.

| Feature | Description | Variable |
| --- | --- | --- |
| *SbjID* | Subject identification number | Ordinal |
| *cosPhase* | Cosine of the phase of auditory stimulation | Continuous |
| *sinPhase* | Sine of the phase of auditory stimulation | Continuous |
| *v.Neg* | Voltage of SO trough before the click | Continuous |
| *v.Stim* | Voltage in the moment of stimulation | Continuous |
| *t.Neg* | Time of duration for the negative wave before click | Continuous |
| *t.PosNeg* | Time of duration for the peak to trough before click | Continuous |
| *t.Stim* | Time between zero-crossing to click time | Continuous |
| *areaPos* | Area under curve for peak section before the click | Continuous |
| *waveRatio* | Duration ratio for the wave before click | Continuous |
| *slopeRatio* | Duration ratio for the negative wave before click | Continuous |
| *FSonStim* | Presence of fast spindle on stimulation (Yes: 1, No: 0) | Binary |
| *SSonStim* | Presence of slow spindle on stimulation (Yes: 1, No: 0) | Binary |
| *FS_Lag* | Time to the previous FS (12 - 16Hz) | Continuous |
| *SS_Lag* | Time to the previous SS (9 - 12Hz) | Continuous |
| *random* | Random dummy variable | Continuous |

**Wave characteristics used as labels:**
| Feature | Description |
| --- | --- |
| $SO_{trough}$ | Trough amplitude of the post stimulus SO |
| $SO_{vpp}$ | Peak to trough amplitude of the SO after the stimulus |
| $SA_{amplitude}$ | Maximal amplitude of the spindle activity locked to the subsequent positive SO after the stimulation |

Two post-stimulus measures were selected for classification of SO analysis and one for SS. All labels to be predicted were computed from wave measurements taken after stimulation. These correspond to the trough amplitude of the post-stimulus SO (SO_*trough*_), the peak-to-peak amplitude of the SO after the stimulus (SO_Vpp_), and the amplitude of the spindle activity locked to the subsequent positive SO after the stimulation (SA_*amplitude*_). The selection of these variables was specified to include one measure with independent dynamics of the ongoing SO (SA_*amplitude*_), one measure with reduced dynamics of the ongoing wave (SO_*trough*_) and one measure including dynamics of the ongoing SO (SO_Vpp_). Furthermore, all of these measure have been shown to be modulated by the CLAS click (Navarrete et al., 2020a; Ngo et al., 2013; Schneider et al., 2020). Nevertheless, as these evaluated measures represent a continuous distribution, post-stimulus measures were binarized for each recording.

Our goal was to investigate whether stimulation results in a strong or a weak neural response. Predicting the post-stimulus values per se using regression would be an interesting alternative approach. However, the physiological function of SO and SS is determined mostly by the global dynamics of these rhythms rather than exclusively their scalp amplitude (Klinzing et al., 2019; Navarrete et al., 2020b). Therefore, an analysis based on predicting strong vs weak response is more informative than the exact prediction of event amplitudes. Thus, we thresholded post-stimulus values into Low and High categories and cast the analysis as a classification problem.

Labelling binarization was performed to determine Low and High values from the selected post-stimulus measures. For this, a thresholding process was applied for each subject. First, outliers’ values were eliminated by removing the highest and lowest 1%. Then, an empirical cumulative function was computed for each recording. For each subject, lower and higher limits were identified independently for each stimulation condition. Values lower than the 25th percentile were marked as the local low threshold, while values higher than the 75th percentile were determined as the high threshold. The middle 50% were then discarded. Subsequently, the low and high thresholds were averaged between conditions for each subject to determine subject-base thresholds across conditions. Then, values higher than the subject-base high threshold were marked as HIGH, and values lower than the subject-base low threshold were marked as LOW. An equal number of LOW and HIGH labels was randomly selected for each subject, and all LOW/HIGH trials of all subjects were used in the training process. Nevertheless, for the testing of each measure we excluded those subjects with a low number of selected trials (N < 10^th^ percentile for all subjects and measures). This led to have a total population of N = 19 for SO_*trough*_ and SO_Vpp_, and a population of N = 16 for SA_*amplitude*_. From this, the SHAM dataset was built with trials identified in the non-stimulation condition whereas the STIM dataset was built with trials marked as auditory stimulated. After pre-processing and trial selection, an average of 132.2(86.6) events remained for each subject, condition, and measure (Total number of trials for SHAM dataset: SO_*trough*_, N = 2890; SO_Vpp_, N = 3036; SA_*amplitude*_, N = 3874. Total number of trials for STIM dataset: SO_*trough*_, N = 1988; SO_Vpp_, N = 1986; SA_*amplitude*_, N = 1946).

The binarization process determined subject-specific threshold values that differentiated low and high amplitude events. Figures 1b-c show the average distribution of labelled LOW and HIGH events. A paired t-test showed that the differences between HIGH and LOW values were significant on a group level for SO_*trough*_ (mean LOW_Thres_ = 24.7uV, 95%CI = 21.7uV to 27.7uV; mean HIGH_Thres_ = 105.6uV, 95%CI = 92.5uV to 118.7uV; t(22.2) = −11.8, p < .001) as well as for SO_Vpp_ (mean LOW_Thres_ = 79.9uV, 95%CI = 73.5uV to 86.3uV; mean HIGH_Thres_ = 188.0uV, 95%CI = 170.5uV to 205.5uV; t(25.3) = −11.4, p < .001) and SA_*amplitude*_ (mean LOW_Thres_ = 3.7uV, 95%CI = 3.5uV to 3.9uV; mean HIGH_Thres_ = 6.5uV, 95%CI = 6.1uV to 6.9uV; t(21.5) = −12.2, p < .001). From these results, we highlight that the LOW_Thres_ for both SO measures are typical for events that are not considered SOs while the HIGH_Thres_ are typical for SOs considered as high amplitude (Iber et al., 2007). The same characteristic can be considered for SA amplitude LOW and HIGH thresholds (Purcell et al., 2017). Hence, it is important to note that the LOW/HIGH values do not differentiate between no-event/event trials, but rather between no-event/high-event trials.

### 2.4 Training of the classifier

Classification models were trained using supervising learning. In supervising learning, these models try to find a good approximation of an unknown function *ƒ* given paired examples (*θ, ƒ(θ)*), where *θ* is a set of features describing each dataset. Several classification algorithms were tested for this study. These included Random Forest (RF), Support Vector Machine (SVM) and Logistic Regression (LR). RF was found to give the highest or close-to-highest accuracies in most cases. SVM was found to give higher accuracies in some cases but the results took significantly longer to compute whereas LR got lower accuracy in the classification. Random forest was therefore chosen because it presented the best trade-off between accuracy and computation time. Likewise, RF has also proved to be efficient in other problems based on sleep EEG (da Silveira et al., 2017; Dimitriadis et al., 2020). Consequently, we independently trained classification models for SHAM and STIM conditions using the RF algorithm.

A RF is an ensemble method or a meta-classifier that combines the outputs of a collection of decision trees. A decision tree is an algorithm that performs classification by performing a cascade of binary decisions (‘branch off left’ or ‘branch off right’). In each binary decision, a single feature is compared against a threshold value. The threshold is determined during training such that the classes are optimally separated using a metric called Gini impurity. A RF creates many such decision trees. To encourage different trees to focus on different aspects of the data, each tree is exposed to a different subset of the data. This is achieved by randomly selecting subsets of the samples. A RF performs classification by collecting ‘votes’ from all the decision trees and then selecting the majority vote. RF has been shown to be a good out-of-the-box classifier that is robust against overfitting, but it is not very sensitive to outliers and its hyperparameters are easy to set. In addition, there is no need for RF to prune the trees which helps in gaining higher accuracy (Breiman, 2001). We implemented decision trees grown with surrogate splits to optimize predictions, achieved by estimating tree splits using complementary features (Hapfelmeier et al., 2012). Hyperparameters were optimized before the learning process. The values were adjusted to optimize the performance of the algorithm. In our RF classifiers, the two hyperparameters were the number of trees in the forest and the maximal number of levels in each decision tree.

For training, all selected trials of all subjects were pooled. To obtain an unbiased estimate of generalisation performance, leave-one-subject-out cross-validation was performed: in each iteration, one subject was left out and the classifier was trained using trials from the other subjects. This was then repeated, leaving each subject out in turn. To tune the model’s hyperparameters, nested cross-validation was performed: in each iteration, the training data was again split into a training set (70%) and a validation set (30%). The validation set was used to find the best hyperparameters. The optimal model was then taken forward and tested on the held-out subject. This resulted in a total of N classifiers being trained for each measure in both SHAM and STIM cases, where N corresponds to the total of subjects evaluated.

Hence, considering *θ_SH_* the set of features for the SHAM dataset, and *θ_ST_* the set of features for the STIM dataset, we will denote as *ƒ_SH_* the RF classifier trained in the SHAM dataset *(θ_SH_, ƒ(θ_SH_))*. Similarly, we will denote as *ƒ_ST_* the RF classifier trained in the STIM dataset *(θ_ST_, ƒ(θ_ST_))*.

To evaluate the performance of the binary classification we used accuracy (ACC) and the Matthews correlation coefficient (MCC) (Boughorbel et al., 2017) which is a robust test for binary decisions. These metrics were calculated according with the predictive forecasting of the binary classification model. Therefore, defining HIGH instances as positive (P) and marked LOW instances as negatives (N), every classification response can be grouped within four cases: i) True positives (TP): positive values that are predicted as positive; ii) False negatives (FN): positive values wrongly marked as negatives; iii) False positives (FP): Negative values classified as positives, and iv) True negatives (TN): actual negative values correctly classified as negatives. Hence, the unbiased performance evaluated in the MCC sense is determined by:

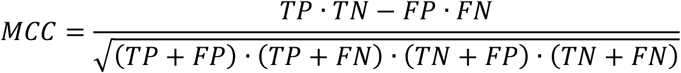

This value ranges from −1 (perfect misclassification) to 1 (perfect classification), where 0 indicates random labelling.

Accuracy accounts for the proportion of correctly classified samples on all classes:

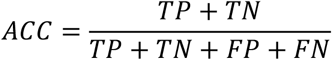

As well as ACC, the MCC gives high scores for correctly labelled classes. Nevertheless, MCC is a stricter measure with high scores only if the model can predict a high percentage of true positives and a high percentage of true negatives on any balanced or imbalanced dataset (Chicco and Jurman, 2020).

Finally, to discern the performance differences in STIM or SHAM datasets, we compared the performance of each model applied within and cross-condition (i.e., SHAM model predicting SHAM labels *ƒ_SH_*(*θ_SH_*) vs STIM model predicting SHAM labels *ƒ_ST_*(*θ_SH_*) as well as SHAM model predicting STIM labels *ƒ_SH_*(*θ_ST_*) vs STIM model predicting STIM labels *ƒ_ST_*(*θ_ST_*).

### 2.5 Feature importance

It is possible to achieve high performance in both *ƒ*_SH_ and *ƒ*_ST_ classifier models. However, the features important for making the prediction may differ between the classifiers, with the importance of some features common to both models. Here we were interested in how the models differed, and which features were uniquely important for one model or the other.

We assessed the relevance of each feature by computing an unbiased measure of feature importance. We implemented a heuristic metric based on the permutation importance of each feature evaluated in the holdout subject (Altmann et al., 2010). For each feature, the algorithm permutes the values of the evaluated feature *S* times and applies the trained algorithm to predict the original labels. The permutation makes a feature uninformative by destroying its relationship with the class labels. The permutation increases the model’s prediction error that derives in a distribution of *S* null performances. The importance of each feature is thus determined by the change in the model’s performance. Hence, big changes in the error are expected for important features whereas none or small changes are expected for less important features.

In our analysis, we first estimated the MCC performance of the original model (*H*). Subsequently, we randomly permute the observations of *x_j_* to estimate the performance of the model using the altered features using one hundred repetitions (*S* = 100) for each feature (*x_j_*, *j* = 1, 2, … N features). For each permuted feature, this resulted in a distribution (*H^*^_j_*) of one hundred performance values. Then, we took the differences *d_j_* = *H* - *H^*^_j_* and computed the mean *D_j_* and standard deviation σ_*j*_. We defined then the feature importance (FI) by permutation for *x_j_* as the z-value *FI_j_* = *D_j_* /σ_*j*_. Changes in performance of the SHAM model in the SHAM dataset (*FI_SH_ = FI(*ƒ_SH_*(θ_SH_))*) and changes of the STIM model in the STIM dataset (*FI_ST_ = FI(ƒ*_ST_*(θ_ST_))*) represent within-condition predictor weights. Likewise, changes in performance of the SHAM model in the STIM dataset (*FI_STx_ = FI(ƒ*_SH_*(θ_ST_))*) and changes of the STIM model in the SHAM dataset (*FI_SHx_ = FI(ƒ*_ST_*(θ_SH_))*) represent cross-condition predictor weights. Finally, we relied on the assumption that highly important features in both models (SHAM and STIM) should be significantly greater than the dummy *random* feature included in the model.

For determining features describing similar dynamics between conditions, we relied on the assumption that features equally important in both models (SHAM and STIM) have similar changes in performance when applied within the same dataset. For each model, the predictor weights represented by the *FI* values represent the vector space of random variables in each dataset. Then, we determined the correlations in between *FI* values when SHAM and STIM were evaluated in the same dataset using the cross-condition models (e.g. FI correlation in SHAM dataset ρ_SH,ST_(*θ_SH_*) = corr(*FI_SH_*, *FI_SHx_*), and FI changes in STIM dataset: ρ_SH,ST_(*θ_ST_*) = corr(*FI_STx_*, *FI_ST_*)). Hence, correlated *FI* variables between models are not independent in the random space (Papoulis and Pillai, 2002), and highly correlated features suggest that these predictors may describe similar processes in SHAM and STIM datasets.

### 2.6 Statistics

To compare classifier performances within and between conditions, we implemented a Wilcoxon signed rank test across holdout subjects. This is equivalent to implementing a 21-fold cross-validated signed-rank test (Thomas, 1998). Also, we evaluated whether the trained models were statistically different by computing the McNemar’s test which evaluates the misclassification rates between models (Thomas, 1998). A one-way analysis of variance (ANOVA) was calculated on feature importance; then the features were compared against the dummy random feature to check for random effects using a Tukey’s Honestly Significant Difference procedure. For comparing EEG trials, we computed significant differences between STIM vs. SHAM within subjects using the Welch unequal variance t-test with the Moser–Stevens correction for degrees of freedom (Moser and Stevens, 1992). The Benjamini & Hochberg false discovery correction (FDR) for multiple comparisons was applied to control for the family-wise error rate (Benjamini and Hochberg, 1995).

## 3. RESULTS

### 3.1 Classification performance

Overall, our trained classifiers achieved good accuracies for both SHAM and STIM models and these were comparable when evaluated on within conditions datasets. Specifically, *ƒ*_SH_ and *ƒ*_ST_ models had comparable performance in all three evaluated measures: SO_*trough*_ (SHAM:ACC= 0.71, 95%CI = 0.69 to 0.74; STIM:ACC= 0.71, 95%CI = 0.67 to 0.74, p = .601), SO_Vpp_ (SHAM:ACC= 0.84, 95%CI = 0.82 to 0.86; STIM:ACC= 0.83, 95%CI = 0.81 to 0.85, p = .295) and SA_*amplitude*_ (SHAM:ACC= 0.60, 95%CI = 0.56 to 0.64; STIM:ACC= 0.59, 95%CI = 0.55 to 0.62, p = .691). This indicates that it is equally possible to predict SO amplitude outcomes from SO morphology in spontaneous and stimulation conditions. However, the amplitude of spindles locked to SOs may be harder to predict as the accuracy was lower when compared to the prediction of SO amplitudes.

We were especially interested in determining whether characteristics of the ongoing spontaneous SO signal could predict if the auditory stimulation would result in a LOW or HIGH SO outcome. To test this, we evaluated the performance of the trained classifier in a cross-classification paradigm. Thus, classifiers trained on the STIM dataset were applied to the SHAM dataset to predict trial labels in SHAM, and classifiers trained on the SHAM dataset were applied to the STIM dataset to predict STIM trial labels. This process helped us to determine the level at which information from unstimulated SWS (SHAM dataset) can be used to predict SO characteristics after an auditory stimulation (STIM dataset), thus providing an idea of how much the auditory stimulation disrupts or alters the ongoing SO pattern. Similar performance of *ƒ*_SH_ and *ƒ*_ST_ when applied within condition indicates a similar degree of generalization for the two models within their own dataset. Equivalent performance of these two classifiers across conditions (cross-classification) would indicate that both classifiers have similar mappings of the feature hyperspace (Thomas, 1998). In other words, similar performance in the cross-condition denotes that the prediction capabilities of these models were mostly based on the characteristics of the spontaneous signal. Conversely, changes in the performance of classifiers in the cross-condition indicate that the mapping of the feature hyperspace is different, and these differences are caused by the auditory click. A Wilcoxon signed rank test revealed no significant difference in within dataset classification for trough, peak-to-peak voltage, or amplitude measures (*ƒ*_SH_(*θ_SH_*) vs *ƒ*_ST_(*θ_ST_*)), for SO_*trough*_ (Z = 0.56, p = .573), SO_Vpp_ (Z = 1.41, p = .159) and SA_*amplitude*_ (Z = 0.62, p = .535) (Figure 2a). Hence, we accept the null hypothesis regarding comparisons of performance of all three measures within the trained datasets suggesting that ƒ_SH_ and ƒ_ST_ perform similarly when applied within condition.

**FIGURE 2.**
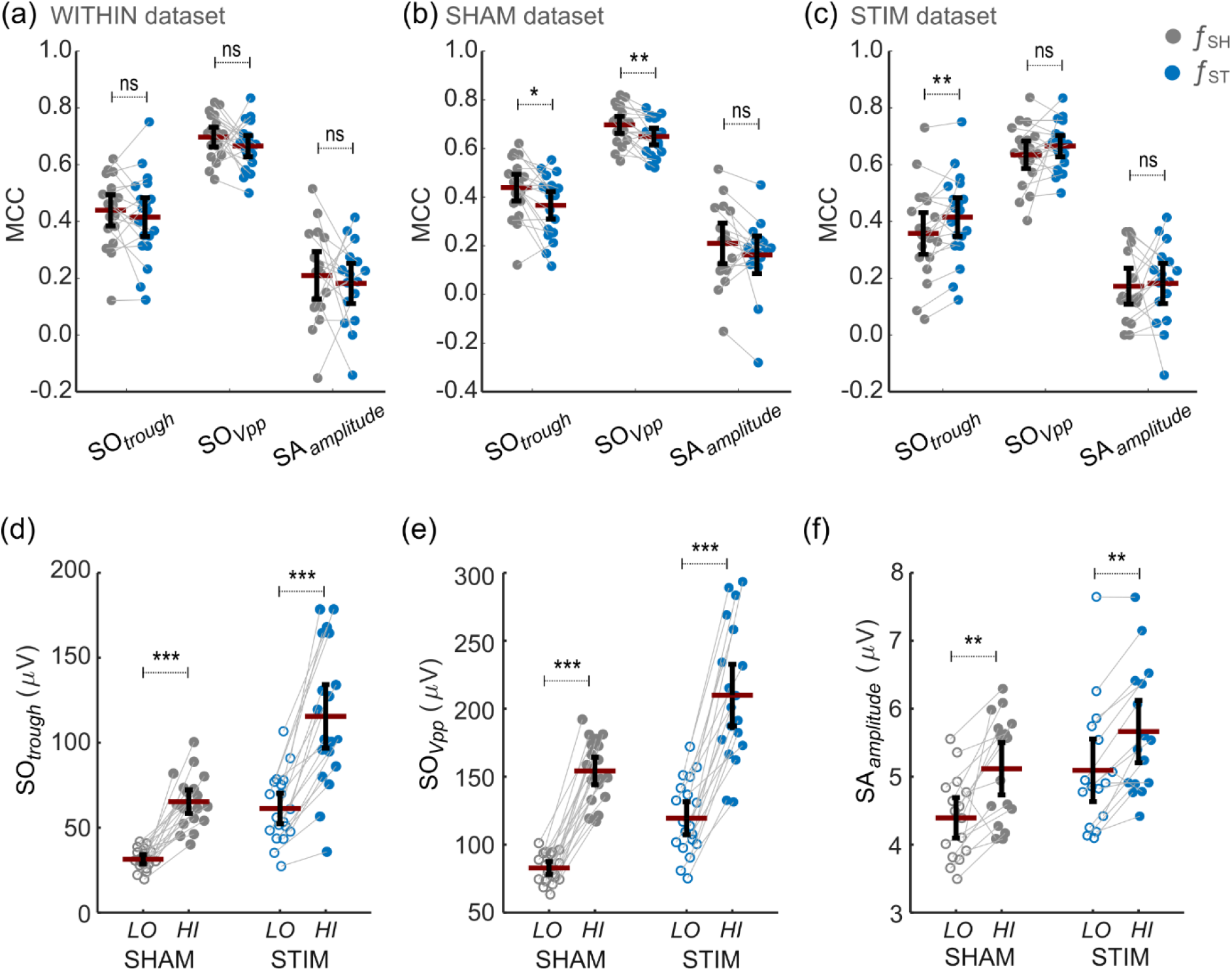
Within and cross-classification performance of trained models and average LOW and HIGH values for post-classification labels. a) Classifier differences in MCC performance for within dataset tests. b) Classifier differences in MCC performance when SHAM classifier (*ƒ*_SH_) and STIM classifier (*ƒ*_ST_) were tested in the SHAM dataset. c) Classifier differences in MCC performance when *ƒ*_SH_ and *ƒ*_ST_ were tested in the STIM dataset. d) Average amplitude of SO_*trough*_ for trials classified as LOW and HIGH in SHAM and STIM datasets. We can notice a larger increment of SO amplitudes in those trials marked as HIGH. The same effect was observed for SO_*Vpp*_ (e) and SA_*amplitude*_ (f). (*) for p < .05, (**) for p < .01, (**) for p < .001, (n.s) for not significant. All error bars represent mean ± 95%CI.

The picture was different in the crossover analysis, where classifiers were trained on one dataset, then applied on both, and classification performance was compared. Firstly, we found significant differences between MCC rates when both classifiers (trained on Stim *ƒ*_ST_ and Sham *ƒ*_SH_) were applied to the SHAM dataset (i.e., *ƒ*_SH_(*θ_SH_*) vs *ƒ*_ST_(*θ_SH_*)). Thus, as shown in Figure 2b, the classification rate in SHAM differed for SO_*trough*_ (Z = 2.55, p = .011) and SO_Vpp_ (Z = 2.77, p = .006), but not SA_*amplitude*_ (Z = 1.34, p = .179). We also found a difference in classification rate when both classifiers were applied to the STIM dataset (i.e., *ƒ*_SH_(*θ_ST_*) vs *ƒ*_ST_(*θ_ST_*)), Figure 3c. Thus, *ƒ*_ST_ outperforms *ƒ*_SH_ when predicting the LOW/HIGH levels of SO_*trough*_ in the stimulation condition (Z = −2.70, p = .007). However, there was no difference in classification performance for SO_Vpp_ (Z = −0.54, p = .586), or SA_*amplitude*_ (Z = −0.57, p = .570).

**FIGURE 3.**
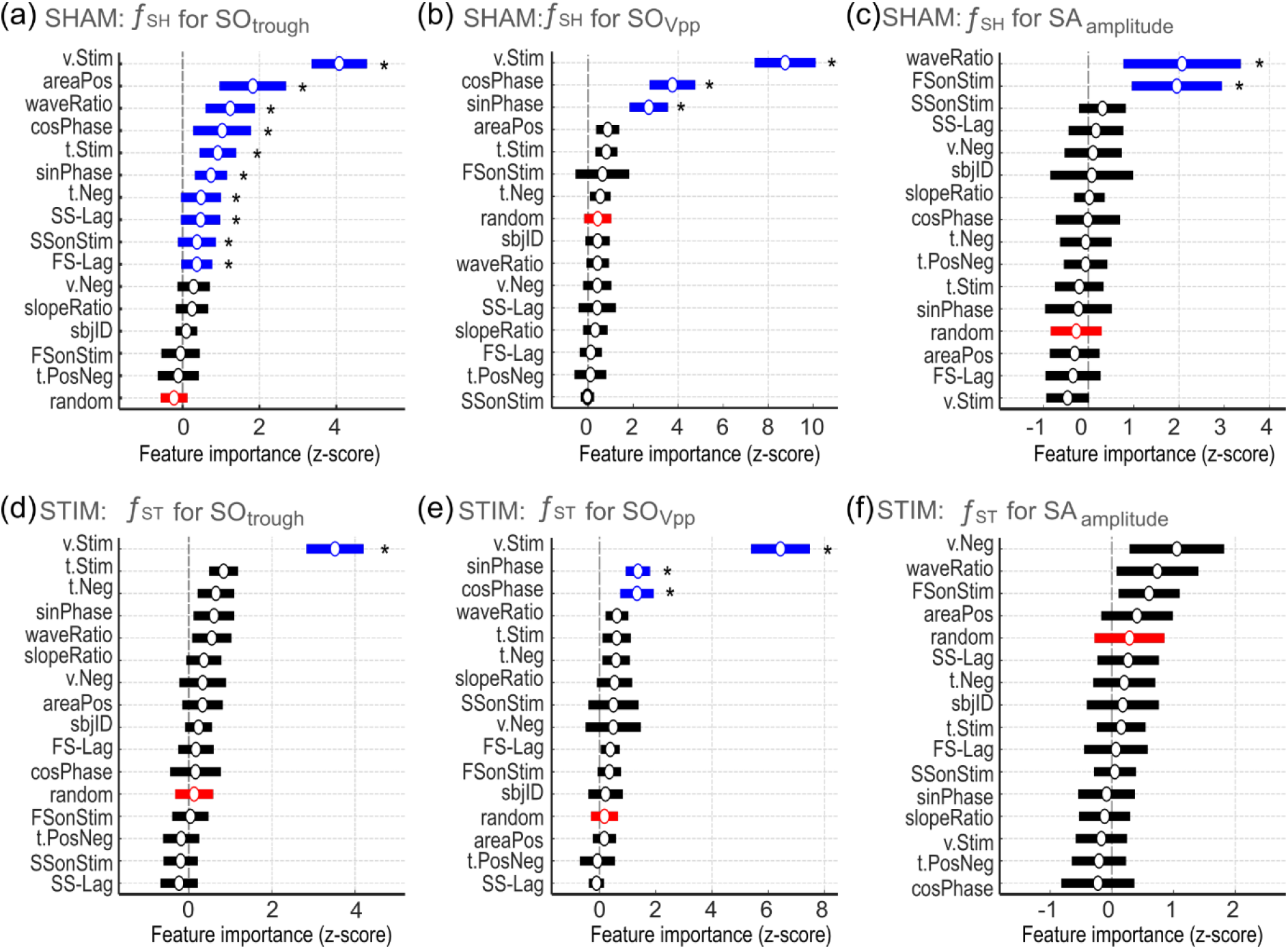
Feature importance of SHAM and STIM classifiers. a) Feature importance by permutation for classification of SO_*trough*_ in the SHAM dataset. b) Feature importance for classification of SO_*Vpp*_ in the SHAM dataset. c) Feature importance for classification of SA_*amplitude*_ in the SHAM dataset. d) Feature importance for classification of SO_*trough*_ in the STIM dataset. e) Feature importance for classification of SO_*Vpp*_ in the STIM dataset. f) Feature importance by permutation (*FI*) for classification of SA_*amplitude*_ in the STIM dataset. Vertical dashed lines at z-score 0 indicate the level of random variability for FI. Feature importance values are provided as the z-score of the permuted MCC performance, and bars in blue indicate those features that are significantly larger than the random dummy feature (in red). (*) for corrected p < .05. All error bars represent mean ± 95%CI

These crossover results suggests that the classifiers (SO_*trough*_; SO_Vpp_; SA_*amplitude*_) differ in how they obtain the information from the data to predict each evaluated measure. Primarily, because we evaluated the performance within condition and found no difference between classifiers, we know that they have similar performance in their respective condition (Fig. 2a). This means that any differences between conditions cannot be attributed to imbalance in the performance of *within* classification. Similarly, looking at the spindle measure, the performance for predicting SA_*amplitude*_ is similar for both classifiers in the crossover analysis (Fig. 2b&c). This indicates that there is not enough information for the *ƒ*_ST_ classifier to predict SA_*amplitude*_ responses beyond the information provided by the spontaneous ongoing SO features in the STIM dataset. Turning to the SO measures (SO_*trough*_ and SO_Vpp_), the SHAM classifier *ƒ*_SH_ outperformed the stimulation classifier *ƒ*_ST_ on both measures in the SHAM dataset *θ_SH_*, (Fig. 2B), but the STIM classifier *ƒ*_ST_ only outperformed the SHAM classifier *ƒ*_SH_ for SO_*trough*_ in the STIM dataset *θ_ST_* vs *θ_SH_*. (Fig.2c). The fact that the STIM classifier did not outperform the SHAM classifier *ƒ*_SH_ for SO_*Vpp*_ in the STIM dataset may indicate that the auditory click does not significantly alter the positive SO amplitude, increasing the complexity of the SO_Vpp_ detection. However, the fact that the STIM classifier outperformed the SHAM classifier on SO_*trough*_ indicates that the pre-stimulus information used to predict SO_*trough*_ is different for the two classifiers. This suggests that the stimulation strongly influences the post-stimulus SO trough, since that trough is no longer predicted by the same pre-stimulus features as it would have been if no click had occurred.

To further corroborate our results, we compared the trained classifiers in each dataset using a more robust method based on their classification errors. Hence, we applied a non-parametric McNemar test which compares the incorrectly labelled trials generated by each classifier. Under the null hypothesis, the two classifiers should have the same error rate, so for instance, the *ƒ*_SH_ and *ƒ*_ST_ classifiers would not be different (Thomas, 1998). After FDR correction, our McNemar’s tests showed that the error rate of SO measures differed significantly between *ƒ*_SH_ and *ƒ*_ST_. This was the case in both the SHAM dataset *ƒ*_SH_(*θ_SH_*) vs *ƒ*_ST_(*θ_SH_*): SO_*trough*_ (p < .001), SO_*Vpp*_ (p < .001); and the STIM dataset *ƒ*_SH_(*θ_ST_*) vs *ƒ*_ST_(*θ_ST_*): SO_*trough*_ (p = .005); SO_*Vpp*_, (p < .001). Furthermore, we did not find differences between classifiers for spindle activity when both the *ƒ*_SH_ and *ƒ*_ST_ were applied in either SHAM (SA_*amplitude*_, p = .308) or STIM datasets (SA_*amplitude*_, p = .138). Overall, these results are perfectly in keeping with the MCC analysis presented above.

Thus, both the analysis of differences in MCC performance and the analysis of differences in the error rate confirm that training classifiers in SHAM and STIM datasets results in two distinct models. This is particularly true for the SO measures (SO_*trough*_ and SO_Vpp_) which learn different dynamics from the evaluated datasets imposed by the auditory stimulus. The lack of forecasting of stimulus-related changes for spindle activity might indicate that the EEG information that predicts these effects is mainly associated with post-stimulus processes.

### 3.2 Generalization of classifier predictions

Next, we wanted to check how accurately our classifiers had predicted whether subsequent SOs would fit the HIGH or LOW amplitude classes by examining the actual amplitudes which occurred. Therefore, we evaluated the response of the classification for all trials. These included trials within LOW and HIGH groups of labels as well as trials falling between the [LOW, HIGH] interval thresholds that were initially discarded during training of the classifiers. The amplitude of post-stimulus responses was analysed with a 2 (Condition: SHAM vs STIM) x 2 (Classification Label: LOW vs HIGH) ANOVA as seen in Figure 2d-f. This showed a main effect of classification label for SO_*trough*_ (F(73,1) = 59.37, p < .001), SO_Vpp_ (F(73,1) = 126.03, p < .001) and SA_*amplitude*_ (F(61,1) = 9.86, p = .003). In all conditions, a Tukey post hoc test revealed that HIGH labels identified higher amplitude events for SO_*trough*_ (SHAM: t(18.0) = −9.1, p < .001; Cohen’s d = 2.88, 95%CI [1.95, 3.82]; STIM: t(18.0) = −8.7, < .001; Cohen’s d = 1.66, 95%CI [0.90, 2.42], Figure 2d), as well as for SO_Vpp_ (SHAM: t(18.0) = −18.3, p < .001; Cohen’s d = 4.04, 95%CI [2.90, 5.18]. STIM: t(18.0) = −12.3, p < .001; Cohen’s d = 2.23, 95%CI [1.40, 3.07], Figure 2e) and SA_*amplitude*_ (SHAM: t(15.0) = −3.0, p = .008; Cohen’s d = 1.03, 95%CI [0.27, 1.79]; STIM: t(15.0) = −3.3, p = .005; Cohen’s d = 0.61, 95%CI [−0.13, 1.34], Figure 2f). We can thus conclude that the classifier accurately predicted whether SOs and SAs will have comparatively large or small amplitudes when it categorises them into LOW or HIGH classes.

### 3.3 Feature importance of evaluated classifiers

The feature importance (FI), as determined for *ƒ*_SH_ and *ƒ*_ST_ classifiers, showed how strongly each pre-stimulus feature weight was when the model was applied to unseen data (holdout subject). Firstly, for each condition and measure, we applied a one-way ANOVA to test whether features differed in importance. Secondly, we identified the features with FI larger than the FI estimated for a dummy random feature using Tukey’s honesty test.

Interestingly, our analysis revealed that permuting features arising from the wave structure and estimated click timing (‘sham-click’) in spontaneous SO dynamics (SHAM condition) caused significant changes in the z-score of MCC (Figures 3a and 3b). We found that the z-scored FI differed between features in the SHAM classifier for SO_*trough*_ (one-way ANOVA, F(15,280) = 14.27, p < .001, Figure 3a). Tukey post hoc tests revealed that several features for which FI was significantly higher than the FI for the dummy random feature (−0.22, 95%CI: −0.57 to 0.12). These included features from the estimated sham-click (*v.Stim* (4.07, 95%CI: 3.35 to 4.79; p < .001), *cosPhase* (1.03, 95%CI: 0.28 to 1.78; p = .008), *t.Stim* ( 0.92, 95%CI: 0.44 to 1.39; p = .001), *sinPhase* (0.74, 95%CI: 0.32 to 1.16; p = .002)) as well as features of the slow-wave structure (*areaPos* (1.83, 95%CI: 0.96 to 2.69; p = .001), *waveRatio* (1.24, 95%CI: 0.60 to 1.88; p = .001), *t.Neg* (0.48, 95%CI: −0.04 to 1.00; p = .030), *SS-Lag* (0.47, 95%CI: −0.04 to 0.98; p = .030), *SSonStim* (0.37, 95%CI: −0.12 to 0.87; p = .047) and *FS-Lag* (0.37, 95%CI: −0.03 to 0.77; p = .030)).

We also found differences of FI between the various features in the SHAM classifier for SO_Vpp_ (one-way ANOVA, F(15,225) = 36.18, p < .001, Figure 3b). A Tukey post hoc test revealed that FI for *v.Stim* (8.75, 95%CI: 7.39 to 10.11; p = .001), *cosPhase* (3.74, 95%CI: 2.72 to 4.76; p < .001), and *sinPhase* (2.69, 95%CI: 1.84 to 3.54; p < .001) were higher than chance and larger than the dummy random feature (0.42, 95%CI: −0.18 to 1.03).

Similarly, we found differences of FI between features in the SHAM classifier for SA_*amplitude*_ (one-way ANOVA, F(15,240) = 4.40, p < .001, Figure 3c). A Tukey post hoc test revealed that *waveRatio* (2.07, 95%CI: 0.77 to 3.36; p = .015) and *FSonStim*(1.95, 95%CI: 0.96 to 2.94; p = .006) were higher than chance and larger than the random dummy feature (−0.27, 95%CI: −0.83 to 0.29).

Interestingly, three features that depend entirely upon the sham-click time (*v.Stim, cosPhase, sinPhase*) have consistently high FI in SHAM. Indeed, the placement of the Sham-click after the detected SO trough is non-random, and the above features were based on this time. Therefore, these features may strongly predict the magnitude of the next oscillation. Specifically, *v.Stim* (amplitude at sham-click time) is highly correlated with the peak amplitude of the sham-stimulated SO (SHAM dataset: *r* = 0.932, *p* < .001). Similarly, *cosPhase* (the cosine of the phase at sham-click time) correlates with the negative-to-positive slope of the SO (SHAM dataset: *r* = 0.207, *p* < .001) and the area of the negative SO deflection before the sham-click (SHAM dataset: *r* = 0.122, *p* < .001). Likewise, *sinPhase* correlates with the negative-to-positive slope of the SO before the sham-click (SHAM dataset: *r* = 0.123, *p* < .001). Overall, these results show that features that seem related only to the timing of the sham-click (i.e. *v.Stim, cosPhase, sinPhase*) also predict characteristics of the SO structure.

In contrast to SHAM, in the STIM dataset the feature importance of only a few variables were significantly higher than the feature importance of the dummy random feature. Thus, FI differed between features for the STIM classifier on SO_*trough*_ (one-way ANOVA, F(15,278) = 12.943, p < .001, Figure 3d). The Tukey post hoc test revealed that *v.Stim* (3.52, 95%CI: 2.83 to 4.20; p < .001) was the only feature higher than the dummy *random* variable (0.13, 95%CI: −0.33 to 0.59). Other click-related FI such as for *tStim* and *sinPhase* were also higher than chance, but they were not statistically different from the dummy *random* variable.

As with the SHAM dataset, the FI differed between features for the STIM classifier on SO_Vpp_ (one-way ANOVA, F(15,241) = 27.958, p < .001, Figure 3e). The Tukey post hoc test revealed *v.Stim* (6.44, 95%CI: 5.39 to 7.48; p < .001), *sinPhase* (1.37, 95%CI: 0.94 to 1.80; p = .004), and *cosPhase* (1.33, 95%CI: 0.73 to 1.93; p = .014) were higher than the *random* dummy feature (0.17, 95%CI: −0.30 to 0.65).

For SA_*amplitude*_, although there was a significant difference between groups as determined by a one-way ANOVA (F(15,239) = 1.805, p = .035, Figure 3f), there were no significant differences between the FI of the *random* dummy variable and the remainder of features.

### 3.4 Similarity of feature importance between SHAM and STIM models

Finally, we wanted to determine whether the features in SHAM and STIM datasets are evaluated in the same way by *ƒ*_SH_ and *ƒ*_ST_ classifiers. We therefore studied how feature permutation changed the classifier performance within and across conditions. Considering that dynamics learnt by *ƒ*_SH_ are based only on the drift of SWA in the SHAM dataset, features evaluated similarly by both classifiers may indicate that these are mostly related to spontaneous EEG activity. From a mathematical perspective, when applied to the same dataset, *ƒ*_SH_ and *ƒ*_ST_ classifiers can be considered as vectorial functions in the same vectorial space. The predictor weights estimated as values of feature importance thus represent the vector space of random variables in the evaluated dataset. Features with correlated feature importance in both *ƒ*_SH_ and *ƒ*_ST_ classifiers are therefore not independent in the multidimensional feature space, suggesting that these predictors may describe similar processes in SHAM and STIM conditions. The correlation of cross-condition changes in performance in the SHAM dataset during the crossover analysis were defined by ρ_SH,ST_(*θ_SH_*) = corr(*FI_SH_*, *FI_SHx_*), while correlations of cross-condition changes in performance in the SHAM dataset were defined by ρ_SH,ST_(*θ_ST_*) = corr(*FI_STx_*, *FI_ST_*).

In Figure 4 we show the cross-correlogram of FI for within vs cross-condition when the classifiers were applied in SHAM and STIM datasets. Our data suggest that the amplitude of the stimulated SO (*v.Stim*) is similar in importance for predicting SO activity in both datasets. After FDR correction, we found the correlations of feature importance were significant for *v.Stim* (within condition) vs *v.Stim* (cross condition) evaluated under SO_*trough*_ (*v.Stim*: ρ_SH,ST_(*θ_SH_*) = 0.92, p < .001 and ρ_SH,ST_(*θ_ST_*) = 0.88, p < .001, Figure 4a and Figure 4b) and SO_Vpp_ (*v.Stim*: ρ_SH,ST_(*θ_SH_*) = 0.92, p < .001 and ρ_SH,ST_(*θ_ST_*) = 0.88, p < .001, not shown). This suggests that data dynamics mapped by this feature in both models may describe similar information in SHAM and STIM datasets. Hence, the *v.Stim* predictor is not an independent variable between spontaneous and stimulation conditions. This shared dependence between conditions suggests that the classifier effect of this feature on the amplitude of the SO outcome is mainly generated from spontaneous neural dynamics rather than dynamics associated with the stimulation.

**FIGURE 4.**
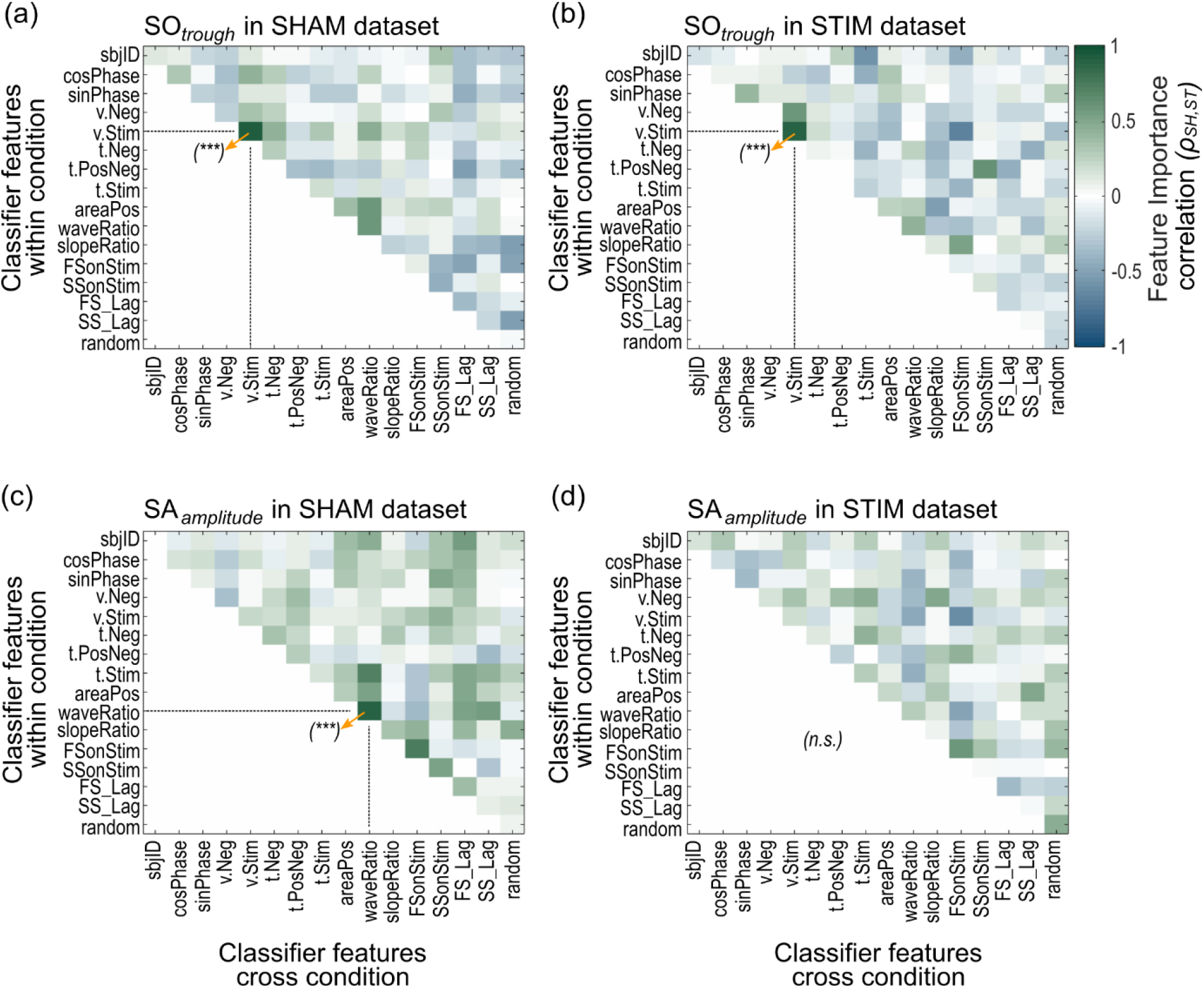
Cross-correlogram for changes in feature importance between within and cross classification models applied to SHAM (ρSH,ST(θSH)) or STIM (ρSH,ST(θ_ST_)) datasets. (a-b) Correlations of feature importance between classifiers applied to SHAM (a) and STIM (b) datasets evaluated on SO_*trough*._ (c-d) Correlations of feature importance between classifiers applied to SHAM (c) and STIM (d) datasets evaluated on SA_*amplitude*_. Significant correlations were highlighted by an arrow and (***) for p < .001 after FDR correction, (n.s) stands for not significant.

Similarly, for SA_*amplitude*_ we found that correlations between feature importance from within and cross condition classifiers were significant for correlation of *waveRatio* (within condition) with *waveRatio* (cross condition) in SHAM (ρ_SH,ST_(*θ_SH_*) = 0.90, p < .001, Figure 4c), but this was not true in the STIM dataset (Figure 4d). This further suggests that the auditory click disturbs the pre-stimulus slow-wave dynamics that predict SA_*amplitude*_. Specifically for the SHAM dataset, we found a high correlation of feature importance between within and cross condition classifiers for *waveRatio* vs *t.Stim* (ρ_SH,ST_(θ_SH_) = 0.72, non-corrected p = .011) and *FSonStim* vs *FSonStim* (ρ_SH,ST_(θ_SH_) = 0.75, non-corrected p = .006) but these did not survive after FDR correction. This further suggests that information from the shape of the SO wave predicts the post oscillation spontaneous spindle activity.

Further electrophysiological analysis of classified trials shows how predicted SO amplitudes (HIGH and LOW classes) mediate the post-click SO or spindle ERP amplitudes in SHAM and STIM datasets. Figure 5a depicts the average ERP for trials predicted as HIGH vs trials predicted as LOW for SO_*trough*_ in the SHAM dataset. Note the apparent increase of SO trough amplitude before the estimated time of stimulation. This could suggest that this trough amplitude (*v.Neg*) is a main predictor for spontaneous SO_*trough*_. However, *v.Neg* was not identified as a top predictor by feature importance as indicated previously in Figure 3a. Hence, bigger troughs do not predict a bigger post-click SOs, but rather the apparent increasing of trough amplitude may correspond to averaging the structure of the negative deflection of SOs, represented by the most important features (*areaPos, waveRatio, cosPhase, t.Stim, sinPhase* as in Figure 3a and section 3.3). As expected, the amplitude of the wave at the estimated click (*v.Stim*) shows larger positive deflections for trials predicted as HIGH SO_*trough*_.

**FIGURE 5.**
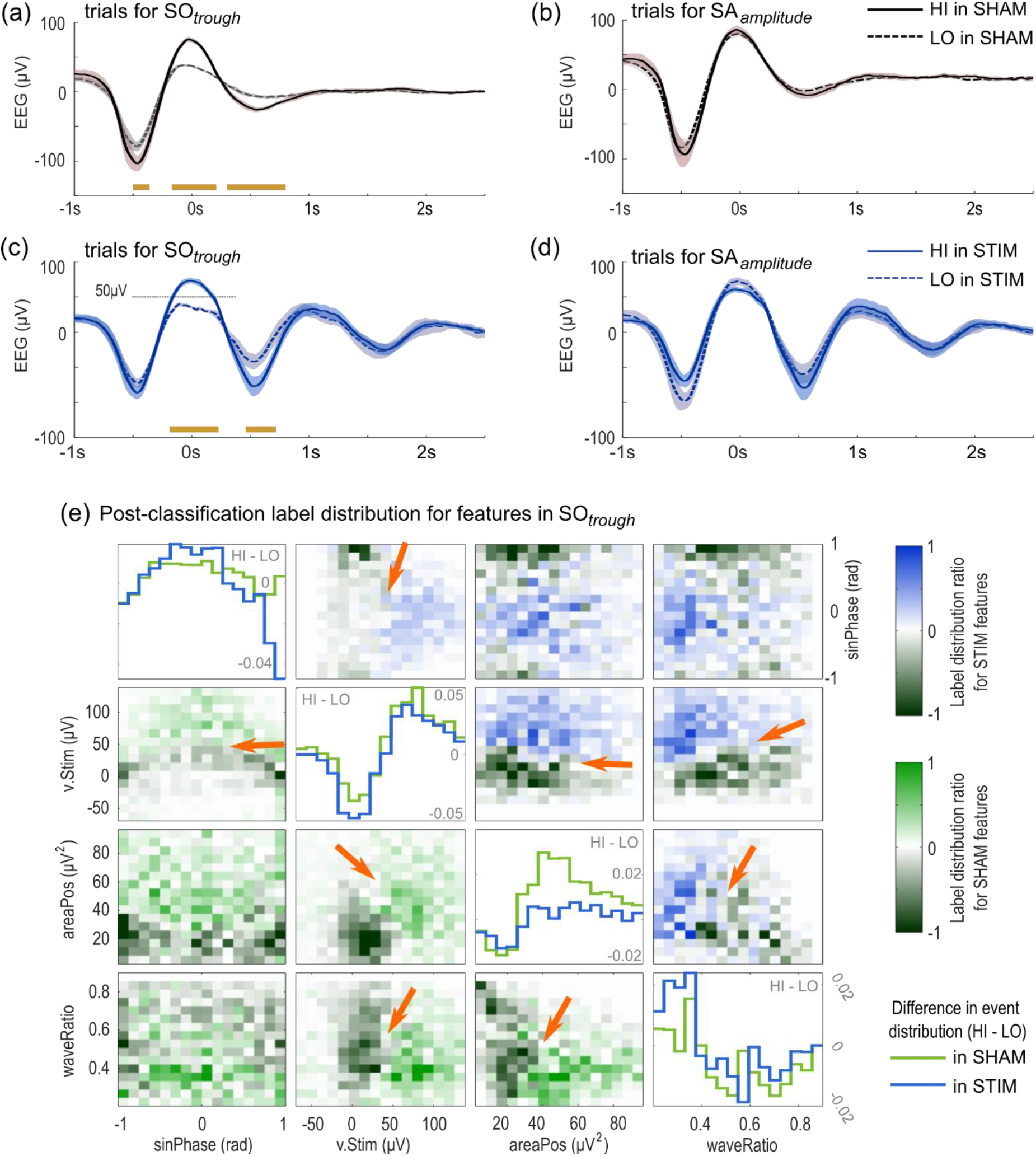
Wave amplitude and feature importance for post-classification labels. (a) Post-classification HIGH (HI) vs LOW (LO) EEG average trials locked to the estimated click for SO_*trough*_ classification evaluated in the SHAM dataset. (b) Post-classification HI vs LO EEG average trials locked to the estimated click for SA_*amplitude*_ classification evaluated in the SHAM dataset. (c) Post-classification HI vs LO EEG average trials locked to the applied click for SO_*trough*_ classification evaluated in the STIM dataset. (d) Post-classification HI vs LO EEG average trials locked to the applied click for SA_*amplitude*_ classification evaluated in the STIM dataset. Orange bars indicate clusters of significant differences between classified LO and HI trials (p < .05) after FDR correction. Error areas represent mean ± 95%CI. (e) Pair plots for some of the top features by FI for SO_*trough*_ indicating the difference of normalized histograms (HIGH - LOW) for SHAM (lower diagonal) and STIM (upper diagonal) datasets. Plots in the diagonal shows the histogram difference between selected features. Positive values indicate larger concentration of HIGH events whereas negative values indicate a larger concentration of LOW values. Orange arrows indicate apparent boundaries between HIGH and LOW events for STIM (black-to-blue) and SHAM (black-to-green) conditions. v.Stim: SO voltage during stimulation; sinPhase: sinus of phase of estimated/applied stimulation; areaPos: Area under curve for the peak of the SO wave before the click, and waveRatio: Duration ratio for the wave before click.

Similarly, Figure 5c depicts the averages of trials predicted as HIGH outcome vs trials predicted as LOW for SO_*trough*_ in the STIM dataset. Unlike predictions in SHAM, in STIM the structure of the pre-stimulus SO did not predict changes in the averaged SO between HIGH and LOW trials. Only the SO amplitude during the click (*v.Stim*) provided a representative feature for SO_*trough*_ prediction.

Conversely, EEG trials representing LOW and HIGH SA_*amplitude*_ were not different for either SHAM (Figure 5b) or STIM datasets (Figure 5d). These findings are in keeping with reduced importance of SO amplitudes for the prediction of post-event SA_*amplitude*_ in both conditions.

Importantly, features should not simply be considered in isolation since they interact together in terms of their impact on the subsequent oscillatory structure. To look at this, we performed a more detailed analysis of the combined contribution of the highest ranked features in the prediction of LOW/HIGH for SO_*trough*_ outcome (Figure 5e). First, to distinguish between the feature distributions, we calculated the difference between normalised event distributions (HIGH - LOW) for SHAM and STIM datasets. The resultant distribution indicates that large *v.Stim* predict larger subsequent SO_*trough*_ amplitudes. Notably, trials having a *v.Stim* > 50 μV are associated with a larger concentration of HIGH events for both STIM and SHAM (Fig 5e). Interestingly, this 50μV threshold is also evident in the averaged SWA in which ERPs were locked to the time of stimulation for HIGH and LOW trials in SHAM (Fig 5a) and STIM (Fig 5c) conditions. Furthermore, the features from the wave structure (*areaPos* and *waveRatio*) also hint at larger effects of these dynamics in the distribution of LOW and HIGH events in both datasets (orange arrows in Fig 5e).

## 4. DISCUSSION

We were able to accurately predict whether application of an auditory click to a SO will lead to a large enhancement of the subsequent SO. This was achieved by using features of an ongoing slow oscillation in a machine learning classifier. Thus, we were able to predict the amplitude of the trough (SO_*trough*_), the peak-to-trough amplitude of the SO (SO_*Vpp*_), and the maximal amplitude of the spindle activity locked to the subsequent positive SO (SA_*amplitude*_) after each click-time in both STIM and SHAM conditions by analysing oscillation features from before the click or sham-click. We also correctly predicted whether either spontaneous oscillatory activity or a phase-locked sound would lead to a large neural response (classification accuracy > 67% for SO amplitude and > 55% for spindle activity) in both STIM and SHAM conditions. Finally, we performed a feature importance analysis to identify the most important EEG features for prediction of SOs and spindles. To this end, we permuted features to assess how they influenced the generalization of classifiers to unseen trials.

### Classification using pre-stim SO wave features differentiates SO click-response in SHAM and STIM

Although our classifiers performed similarly in SHAM and STIM datasets, performance was decreased in cross-classification for both datasets. This is likely due to disruption in the ongoing EEG pattern caused by the auditory click, since the click is the only thing that distinguishes the two conditions. This pattern revealed that STIM and SHAM classifiers represent different mathematical models that derive different information from the ongoing signal in these two conditions, at least when classifying for SO_*trough*_ and SO_*Vpp*_. These results demonstrate that the EEG features prior to the presentation of the sound stimulus carry information about the magnitude of the subsequent brain response following the sound. Conversely, we found no differences between classifiers for SHAM and STIM when predicting SA_*amplitude*_, possibly indicating that pre-stimulus information is not very helpful when predicting this measure in the STIM dataset.

### Spontaneous pre-stim features predict post-stimulus SHAM and STIM spindle dynamics

Overall, we found no differences in the model prediction of spindle activity amplitude between SHAM and STIM models. Previous studies indicated a boosting effect of CLAS on spindle activity. This enhancing effect is specifically related to an increase in amplitude of spindle activity locked to the SO directly after the click (Ngo et al., 2013; Schneider et al., 2020). We found that the presence of spindle events in the ongoing wave and the SO structure were the main variables predicting spindle amplitude. Furthermore, we found evidence that CLAS disturbed the spontaneous relationship between post-stimulus spindle activity and ongoing SO dynamics. However, the ongoing SWA appears to retain the same level of information for SHAM and STIM classifiers to predict the amplitude of post-stimulus spindle activity. Consistent with previous literature (Antony et al., 2019; Ngo et al., 2015), we found that changes in spindle related activity (e.g. FSonStim) were better predicted by refractory periods of spindle rebound. Our results further support the idea that responses to external stimuli such as clicks diverge for innate SWA because of the distinctive thalamocortical dynamics of SO and spindles (Navarrete et al., 2020a). Hence, the pre-stimulus conditions evaluated here in both STIM and SHAM models only conveyed information relating to spontaneous spindle dynamics, and information relevant to the spindle response to CLAS might be mainly concealed within post-stimulus SWS dynamics.

### Feature importance in classification of the SHAM dataset

Analysis of feature importance for the SHAM classifier allowed us to determine which features best characterize the level of cortical activity of the SO cycle. We found that variables relating to the timed structure of the SO were important for predicting spontaneous SO amplitudes. Indeed, because the timing of stimulation depends on the wave detection algorithm (Navarrete et al., 2020a; Ngo et al., 2015), the stimulation-related variables were given high feature importance accordingly. Our results thus suggest that sham-click stimulation related features in the SHAM dataset also convey information about subsequent spontaneous cortical activation. Specifically, the voltage during the sham-click (*v.Stim*) may indirectly reveal the level of spontaneous neural depolarization, which is then measured via scalp amplitude (Crunelli et al., 2018; Siclari et al., 2014). Meanwhile, global neural synchrony may be determined by the timing and phase of the estimated stimulus (*cosPhase, sinPhase* and *t.Stim*) along with the information about SO wave structure (*areaPos, waveRatio, slopeRatio*, etc) (Riedner et al., 2007). Therefore, the features we identified as important for the prediction of spontaneous SO activity (amplitude: *v.Stim*; and SO structure: *areaPos, waveRatio, t.Neg, cosPhase, sinPhase*) may also index the strength of thalamocortical drive on individual SOs.

Following this, we hypothesize that the prediction of spontaneous SO activity by the SHAM classifier may help to discriminate thalamocortical from cortico-cortical SOs. Recent studies proposed that synchronizing factors of the SO may assist in discerning different thalamic and cortical mechanisms involving the dynamics of the ongoing SWA (Bernardi et al., 2018; Siclari et al., 2014). In this direction, thalamocortical circuits trigger Up-states on each SO wave, contributing to cortical synchronization across the cortex and driving the structure of the SO cycle (Amzica and Steriade, 1995; Crunelli et al., 2018) and thus may be indicated by those features describing the SO structure. Likewise, the high importance of the scalp amplitude predictor (*v.Stim*) agrees with previous work indicating that increased levels of cortical depolarization may lead to large synchronous hyperpolarisations (Neske, 2016), and therefore this could highlight intrinsic characteristics of the thalamocortical dynamics (Siclari et al., 2014). Nevertheless, this hypothesis should be tested by adding a larger spatial sample and further intracortical recordings.

### Feature importance in the STIM classifier indexes SO dynamics modulating response to the auditory click

Our feature importance analyses for the STIM classifier indicated that the main variable predicting the post-stimulus SO-amplitude outcome is the wave amplitude during stimulation. This result follows previous work which has suggested that the peak amplitude is the optimal timing for the auditory click to enhance the ongoing waves (Ngo et al., 2015, 2013). Our findings build on this by indicating that considering the stimulus phase alone is not enough to optimally enhance the SO. Instead, the wave amplitude during stimulation better predicts the outcome of the acoustic click. Consistent with this, we recently demonstrated that the window of opportunity for the click to effectively enhance SOs is limited to a wide phase interval around the wave peak (Navarrete et al., 2020a), challenging the hypothesis of a particular SO phase as the optimal target for SO enhancement (Santostasi et al., 2015). Neglecting the importance of wave amplitude could explain the failure of many auditory clicks to boost SO memory consolidation function while still increasing the post-stimulus response (Henin et al., 2019). In the current report, we show that the average amplitude of post-stimulus trials increases considerably when our classification method is used to select the SO cycles to stimulate (Cohen’s d > 1.6 for SO amplitude and Cohen’s d > 0.6 for spindle activity). We therefore suggest that future detection algorithms may consider inclusion of decision rules that evaluate amplitude characteristics to boost the effectiveness of the acoustic click for enhancing the SO response.

Our results also suggest that auditory clicks during elevated neural excitability within SO peaks set the arousal threshold which contributes to the cortical synchronization process of slow-waves. The predictor importance analyses for the STIM classifier showed that stimulation disturbs the effect of the SO wave structure on the prediction of SO troughs. However, wave amplitude during the click still drives the click outcome as it does in the spontaneous generation of large SO troughs in SHAM. Consistent with this, previous studies have indicated that cortical excitability is maximal during periods of neural activation within SO peaks (Massimini, 2002; Rosanova and Timofeev, 2005). Likewise, it has been proposed that the diffusely projected “matrix” thalamic system imposes a sensory threshold upon the arousal system, allowing auditory stimulation to effectively enhance or decrease SOs, or provoke cortical arousals (Bellesi et al., 2014). We suggest that this arousal threshold is determined by the amplitude of the SO. This could be partly mediated by thalamocortical activity, since this drives synchronous firing in the cortex (Neske, 2016; Siclari et al., 2014). Importantly, cortical activity also suppresses the locus coeruleus such that it is harder for it to trigger arousal during large cortical up-states (Bellesi et al., 2014; Eschenko et al., 2012). Consequently, because of the increased excitability of the cortical up-states in SO peaks and the reduced cortico-coerulear interactions during large SO peaks, the stimulus during high amplitude waves induces a global hyperpolarizing effect which manifests in large SO troughs.

### Study limitations

We would also like to caution the reader about some particularities of our study. For instance, our definition of SA amplitude is broader than the definition of spindle events as we did not evaluate detected spindles. Nevertheless, SA amplitude and spindle detection are closely entwined. Likewise, we did not rule out the possibility that other SO or spindle features may also contribute to the prediction of spontaneous or boosted SO oscillations. Nevertheless, in this study we included the most prominent morphological features of the SO wave, and those that are less likely to be perturbed by the auditory click while preventing feature multicollinearity.

### Conclusions

To summarize, we trained machine learning algorithms to predict spontaneous and post-stimulus SO and SA amplitudes. We found that spontaneously generated SO trough amplitude can be predicted by the ongoing structure of the previous SO wave. By contrast, stimulus-related increase in a SO is mainly predicted by SO amplitude at the time of the click. Therefore, we suggest that SO amplitude may work as a kind of cortical threshold to prevent the click from causing arousal, while maintaining maximal cortical activity.

Based on these findings, we suggest that the online detection of a few salient features and the application of our random forest classifier could be used in the optimization of future CLAS algorithms. Using this method, future studies could evaluate the effect of less stimuli but improved SO response on memory and sleep architecture. Applying optimized version of CLAS may also facilitate a better understanding of the dynamics of both spontaneous SOs and SOs that have been boosted through CLAS.

## CREDIT AUTHOR STATEMENTS

**Miguel Navarrete:** Conceptualization, Methodology, Software, Validation, Formal Analysis, Investigation, Resources, Data Curation, Writing-Original Draft, Writing - Review & Editing, Visualization.

**Steven Arthur:** Conceptualization, Methodology, Software, Validation, Formal Analysis, Investigation, Resources, Data Curation, Writing-Original Draft, Writing - Review & Editing, Visualization.

**Matthias Treder:** Conceptualization, Methodology, Investigation, Writing - Review & Editing, Supervision.

**Penelope A. Lewis:** Conceptualization, Methodology, Resources, Writing - Review & Editing, Supervision, Project administration, Funding acquisition.

## ACKNOWLEDGEMENTS

MN and PL were supported by the European Research Council (681607), SA was supported by a generous donation from Mr. Bernard Cheek. The authors would like to thank L. Satamaria-Comarruvias and M. Rakowska for their helpful comments and discussion on this manuscript.

## DATA AND CODE AVAILABILITY

The data used in the study as well as the software and scripts used to compute the analysis have been made publicly available via the Open Science Framework and can be accessed at: https://osf.io/j9vka/

